# *Helicobacter hepaticus* as disease driver in a novel CD40-mediated model of colitis

**DOI:** 10.1101/2020.04.21.053066

**Authors:** Verena Friedrich, Ignasi Forne, Dana Matzek, Diana Ring, Bastian Popper, Lara Jochum, Stefanie Spriewald, Tobias Straub, Axel Imhof, Anne Krug, Bärbel Stecher, Thomas Brocker

## Abstract

Gut microbiota and the immune system are in constant exchange, which shapes both, host immunity and microbial communities. Here, improper immune regulation can cause inflammatory bowel disease (IBD) and colitis. Antibody therapies blocking signaling through the CD40 – CD40L axis showed promising results as these molecules have been described to be deregulated in certain IBD patients. To better understand the mechanism, we used transgenic DC-LMP1/CD40 animals, which lack intestinal CD103^+^ dendritic cells (DCs) and therefore cannot induce regulatory T (iTreg) cells due to a constitutive CD40-signal in CD11c^+^ cells. These mice rapidly develop spontaneous fatal colitis with an increase of inflammatory IL-17^+^IFN-*γ*^+^ Th17/Th1 and IFN-*γ*^+^ Th1 cells. In the present study we analyzed the impact of the microbiota on disease development and detected elevated IgA- and IgG-levels in sera from DC-LMP1/CD40 animals. Their serum antibodies specifically bound intestinal bacteria and we identified a 60 kDa chaperonin GroEL (Hsp60) from *Helicobacter hepaticus* (*Hh*) as the main specific antigen targeted in absence of iTregs. When rederived to a different *Hh*-free SPF-microbiota, mice showed few signs of disease without fatalities, but upon recolonization of mice with *Hh* we found rapid disease onset and the generation of inflammatory Th17/Th1 and Th1 cells in the colon. Thus, the present work identifies a major bacterial antigen and highlights the impact of specific microorganisms on modulating the host immune response and its role on disease onset, progression and outcome in this colitis model.

## Introduction

The large intestine is colonized with about 10^11^ - 10^12^ bacterial cells / g of luminal content ^1^, to which mucosal immune cells are constantly exposed. These interactions are indispensable to generate tolerance towards harmless commensals or immunity to invading pathogens. It is commonly accepted that the intestinal microbiota has a critical impact on modulating host immune responses in both health and disease ^2, 3, 4^. However, multiple genetic and environmental factors such as immune deficiency, infection, inflammation or antibiotic treatment can alter the microbial composition and direct mucosal homeostasis towards dysbiosis. Inflammatory Bowel Disease (IBD) is linked to dysbiosis and many studies could reveal altered bacterial compositions in IBD patients ^5, 6, 7^. However, it still remains elusive whether dysbiosis is the cause or rather a consequence of IBD ^8^.

IL-17-producing T helper (Th17) cells are not detectable in the intestine of germ-free mice but can be effectively induced in the small intestinal Lamina Propria (LP) by mono-colonization with segmented filamentous bacteria ^9, 10^. Also, regulatory T cells (Tregs) are known to be affected by the gut microbiota as Clostridium clusters IV and XIVa are potent drivers of IL-10^+^Helios^−^ induced T regs (iTregs) ^11^. Furthermore, the human commensal *Bacteroides fragilis* is capable of promoting mucosal tolerance as its polysaccharide A leads to differentiation of CD4^+^ T cells into IL-10 producing Tregs in the steady state but also under inflammatory conditions ^12^. Of note, *Bacteroides* can also contribute to disease development under certain conditions ^13^ and, *B. fragilis* was reported to be enriched in IBD patients ^14^. These examples and other reports illustrate how the immune system is shaped by microbiota of the gut. Similarly, the murine commensal *Helicobacter hepaticus* (*Hh*) is found in many academic and commercial mouse colonies ^15, 16^ and infection with *Hh* is linked to chronic hepatitis as well as hepatocellular carcinoma ^17, 18^. *Hh* is also able to elicit intestinal inflammation in immunodeficient or immunocompromized mice. For example, adoptive transfer of CD4^+^ T cells into mice with severe combined immunodeficiency (scid) ^19^ or Rag2-deficiency ^20^ develop colitis only in presence of *Hh*. Also, IL-10-or T cell-deficient mice require *Hh* for development of IBD ^21 22, 23^. However, there are still major gaps in our understanding of the complex interaction between the microbial community and/or certain single species and the host.

We recently published a novel CD40-mediated mouse model of spontaneous colitis, where CD11c-specific constitutive CD40-signaling leads to migration of CD103^+^ DCs from the colonic LP to draining lymph nodes followed by DC-apoptosis ^24^. Loss of tolerogenic CD103^+^ DCs caused a lack of RORγt^+^Helios^−^ iTregs and an increase of inflammatory IL-17^+^IFN-γ^+^ Th17/Th1 and IFN-γ^+^ Th1 cells in the colon, resulting in the breakdown of mucosal tolerance and fatal colitis ^24, 25^. A consequence of this IBD was malabsorption of nutrients and cholesterin due to IBD ^26^. Of note, this model mimics the human IBD situation, as CD40-CD40L interactions are of relevance to the pathogenesis of IBD ^27, 28, 29, 30, 31, 32, 33^.

In the present study we focused on microbial-host interactions in the CD40-mediated colitis model to determine how the intestinal microbiota can modulate the host immune response. We identified *Hh* as disease driver with impact on disease onset, progression and outcome in mice with DC-specific constitutive CD40-signaling. The immune response of diseased animals targets *Hh* and we identified GroEL, a 60kDa *Hh*-protein as a main antigen recognized by immunoglobulins during onset of fatal IBD. Rederivation of the mice to *Hh*-free state saved mice from fatal colitis. This suggested, that *Hh* could trigger colitis and specific immune responses in an iTreg-free setting.

## Results

### Early disease onset in DC-LMP1/CD40 mice is associated with increasing serum antibody levels specific for bacterial antigens

To obtain further insight into the complex interplay of microbiota, adaptive immunity and inflammation in CD40-mediated colitis, we first determined the disease onset in DC-LMP1/CD40 mice by measuring fecal lipocalin-2, a sensitive non-invasive inflammatory marker ^34^. Lipocalin-2 levels were significantly increased in DC-LMP1/CD40 mice starting on week 5 (Fig. 1A), indicating a very early disease onset due to constitutive CD40-signaling on DCs as published previously ^24^. To measure a potential impact on adaptive immunity, we then analyzed IgG and IgA serum levels in these animals during colitis progression. Compared to control littermates, DC-LMP1/CD40 mice showed elevated total serum IgG-as well as IgA-levels already at 6 weeks and increased further with age (Fig. 1B). As mice with spontaneous colitis have the propensity to develop antibody responses against commensal bacteria ^35^, we next set out to identify antibody specificities in DC-LMP1/CD40 mice. To this end we used cecal bacterial lysate (CBL) from non-transgenic C57BL/6 mice of the same colony, representing unaltered intestinal microbiota for ELISA ^35^. In DC-LMP1/CD40 mice serum IgG response to commensal antigens was significantly increased at the 10-week time point if compared to control littermates (Fig. 1C, left). In contrast, we detected significantly higher serum IgA reactivities starting at the age of 10 weeks at all time points analyzed (Fig. 1C, right). To further visualize bacterial antigens potentially recognized by serum Ig from DC-LMP1/CD40 mice, we tested these sera also by immunoblotting (Fig. 1D). Serum IgG from both, DC-LMP1/CD40 mice and control littermates, detected some proteins of different sizes ranging from 10 to 250 kDa (Fig. 1D, left). However, in contrast to sera from controls, each serum IgG sample from DC-LMP1/CD40 mice showed reactivity with a protein of about 60 kDa (Fig. 1D, left). This reactivity increased with the age of mice (Fig. 1D, left). Also, serum IgA from 10-, 12- and 14-week samples of DC-LMP1/CD40, but not control mice, detected proteins around 60 kDa (Fig. 1D, right). Our data reveal a very early disease onset in DC-LMP1/CD40 mice simultaneously with an increase of serum reactivity against commensal antigens present in CBL from healthy mice of the same colony.

**Figure 1:**
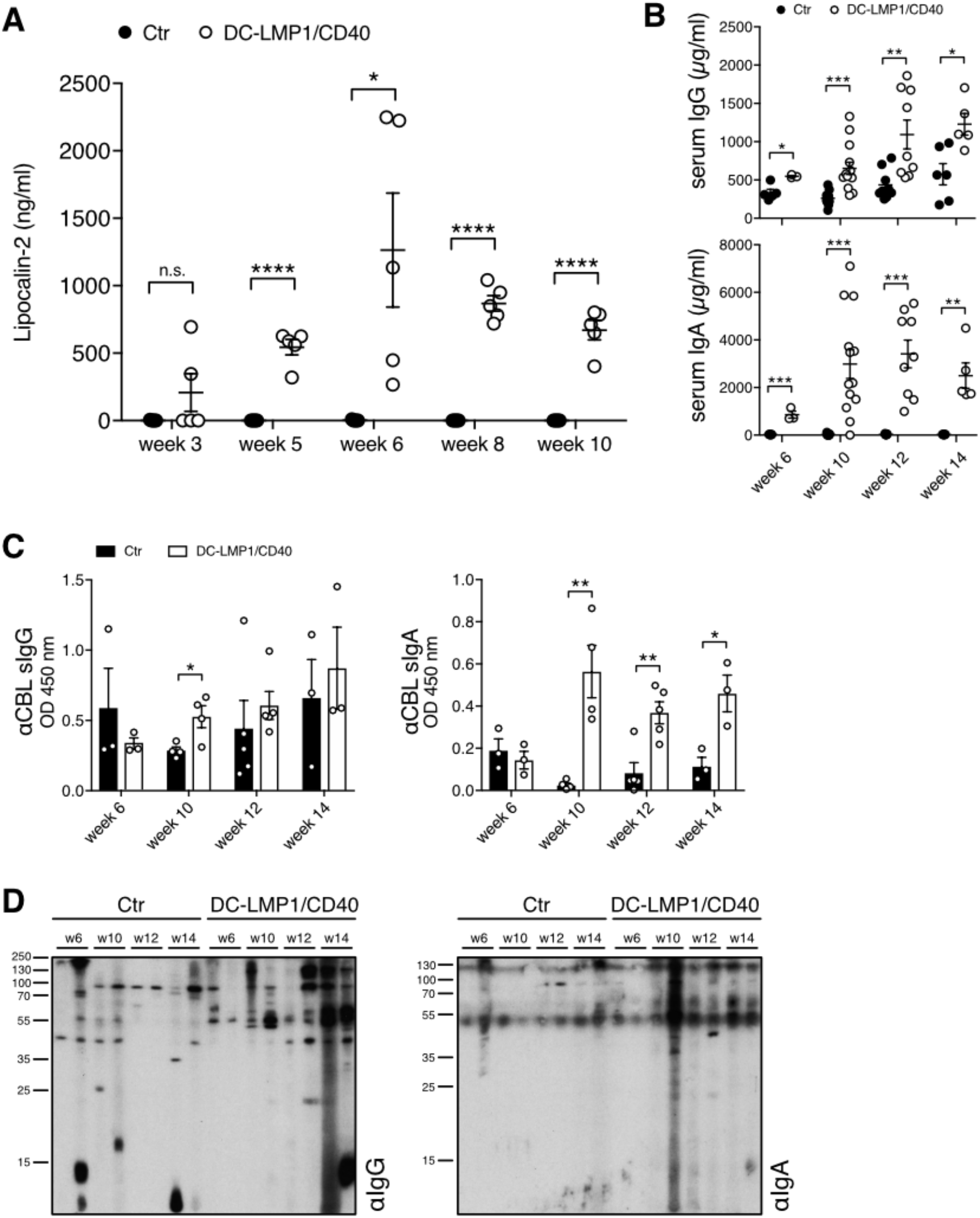
Early disease onset and increasing serum antibody titers. (A) Levels of fecal lipocalin-2 were measured by ELISA in Ctr and DC-LMP1/CD40 mice at indicated time points. Data is shown as mean ± SEM (n=5). (B) Total IgG (upper panel) or IgA (lower panel) concentrations in sera from Ctr and DC-LMP1/CD40 mice at the indicated time points were measured by ELISA. Data from two pooled experiments is shown as mean ± SEM (n=3-13). (C-D) Serum IgG (left) and IgA (right) response in Ctr and DC-LMP1/CD40 mice towards commensal antigens within the CBL was determined by (C) ELISA (mean ± SEM, n=3-5 per group and time point) or (D) immunoblotting at the indicated time points (n=2 per group and time point, each lane represents one serum sample from Ctr or DC-LMP1/CD40 mice). Goat anti-mouse IgG-HRP or goat anti-mouse IgA-HRP were used as secondary antibodies.

### Serum antibodies from DC-LMP1/CD40 mice are specific for a 60 kDa chaperonin from *Helicobacter hepaticus*

To identify antigens recognized by serum Ig in DC-LMP1/CD40 animals, we performed liquid chromatography tandem mass spectrometry (LC-MS/MS) (Fig. 2A). For this approach, serum antibodies were coupled to beads and incubated with CBL for binding of potential target proteins. Upon immunoprecipitation we performed on-bead digestion of proteins followed by LC-MS/MS. The resulting peak intensities were finally used for intensity-based absolute quantification (iBAQ). Proteins identified with a fold change > 2 and a *p*-value < 0.05 were considered for further analyses. Interestingly, the results provided only five proteins precipitated by serum antibodies from DC-LMP1/CD40 mice and two proteins by control serum antibodies (Fig. 2B) that met these requirements. We focused on proteins precipitated by serum antibodies from DC-LMP1/CD40 animals with the highest fold change and lowest *p*-value, which were (i) the 60 kDa chaperonin GroEL (Hsp60) from *Helicobacter hepaticus* (*Hh*) (CH60_HELHP, 8.36-fold change, *p*-value < 0.00001) and (ii) the probable peroxiredoxin from *Helicobacter pylori* (TSAA_HELPJ, 9.93-fold change, *p*-value < 0.000001). The data analysis for the number of precipitated peptides and the percentage of sequence coverage of the protein revealed that the CH60_HELHP was identified by 1-21 peptides with a sequence coverage ranging from 2.4 % up to 43.7% (Fig. 2C). In contrast, TSAA_HELPJ was identified by only one peptide and with a sequence coverage of only 5.6 % for every single DC-LMP1/CD40 serum sample (Fig. 2C). This protein from *H. pylori* was not considered for further analyses as both, the numbers of peptides as well as the percentage of protein sequence coverage were not reliable. One explanation for recovering a protein from *H. pylori* with this approach might be the fact that about 50 % of total proteins from *Hh* have orthologs in *H. pylori* ^36^ and therefore might arise by the analysis within the bacterial database used for iBAQ. Indeed, blasting the precipitated *H. pylori* peptide against the *Hh* proteome resulted in 100 % identity with peroxiredoxin from *Helicobacter* multispecies as well as 70 % identity with chemotaxis protein from *Hh*.

**Figure 2:**
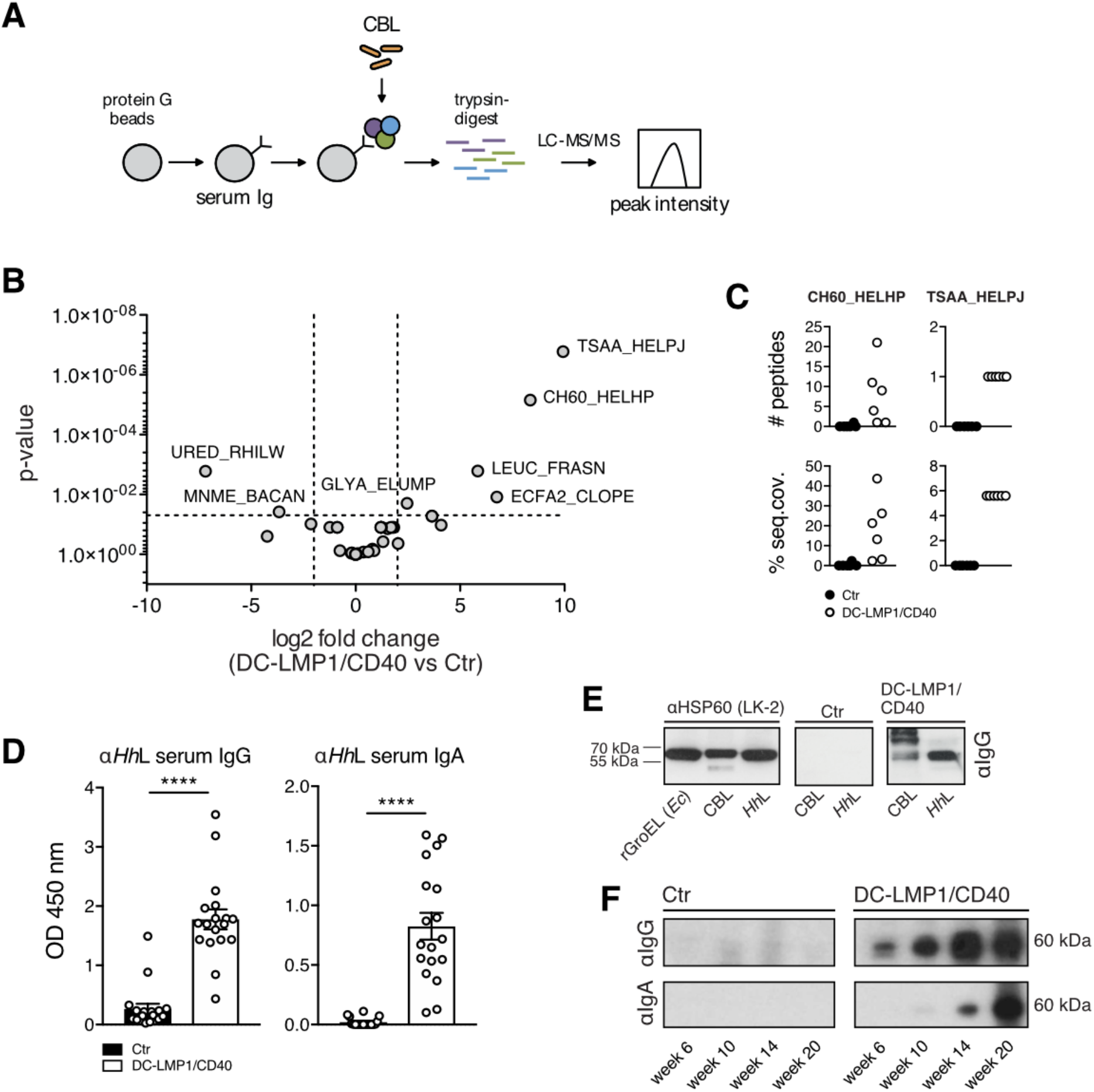
Analysis of fecal antigens. (A) Schematic illustration of sample preparation for liquid chromatography tandem mass spectrometry (LC-MS/MS). Protein G beads were coupled with serum antibodies from Ctr or DC-LMP1/CD40 mice to bind commensal antigens within the CBL. Upon immunoprecipitation, proteins were trypsin-digested, analyzed by LC-MS/MS and the resulting peak intensity was used for intensity-based absolute quantification (iBAQ) (pooled results from two experiments, n=6). (B) Results obtained with iBAQ as described in (A) are illustrated by the volcano plot. Identified proteins were considered as interaction partners if the log2 difference between the iBAQ values in the DC-LMP1/CD40 condition and the controls were higher than 2 and the p-value smaller than 0.05 (ANOVA). (C) Data illustrates the number of peptides (upper panel) and percentage of protein sequence coverage (lower panel) of the identified CH60_HELHP and TSAA_HELPJ in (B). Each symbol represents one single mouse. (D) Serum IgG (left) and IgA (right) response in Ctr and DC-LMP1/CD40 mice towards lysate from *Hh* (*Hh*L) was determined by ELISA (mean ± SEM, n=18). (E) Detection of the 60 kDa protein in *Hh*L and CBL by immunoblotting. 20 μg *Hh*L, 50 μg CBL or 0.5 μg recombinant GroEL from *E.coli* (rGroEL (*Ec*)) were separated by SDS-PAGE. Anti-HSP60 (left, clone LK-2: recognizing both human and bacterial Hsp60, mouse IgG1 isotype) as well as sera from Ctr (middle) and DC-LMP1/CD40 (right) mice were used as primary antibodies. Anti-mouse IgG-HRP was used as secondary antibody. (F) Sera screening for the detection of the 60 kDa chaperonin from *Hh* by immunoblotting. 200 μg *Hh*L were separated by SDS-PAGE and sera from Ctr or DC-LMP1/CD40 mice at the indicated age were used as primary antibodies with each lane representing one serum sample. Anti-mouse IgG-HRP (upper panel) or anti-mouse IgA-HRP (lower panel) were used as secondary antibodies.

To exclude biased results due to differences in serum antibody amounts from DC-LMP1/CD40 and control animals bound by protein G beads, samples were adjusted by calculating equal amounts of serum IgG before coupling onto the beads and also the peak intensities of Ig-related proteins were quantified within the same experiment. Here, DC-LMP1/CD40 and control serum samples showed no differences in Ig-related protein intensities (Fig. S1), indicating equal coupling of serum Ig from control and transgenic mice.

We next tested serum antibody reactivity from DC-LMP1/CD40 mice towards whole *Hh*-lysate (*Hh*L) by ELISA (Fig. 2D) and immunoblotting (Fig. 2E, 2F). Indeed, both, serum IgG as well as IgA from DC-LMP1/CD40 mice showed a strong reactivity towards *Hh*L when compared to sera from control littermates by ELISA (Fig. 2D). To detect GroEL from *Hh* by immunoblotting, we used the monoclonal anti-human heat shock protein 60 (αHsp60) antibody (clone LK-2, mouse IgG1 isotype) as positive control, which specifically recognizes both, human Hsp60 and the bacterial homologue GroEL ^37^. As expected, in Western blot analyses αHsp60 (LK-2) detected recombinant GroEL from *E. coli* (rGroEL (*Ec*)), from CBL and from *Hh*L (Fig. 2E, left panel), confirming the specificity of this Ab and the presence of GroEL in CBL used for this screening. Furthermore, in contrast to sera from control littermates (Fig. 2E, middle panel), sera from DC-LMP1/CD40 mice (Fig. 2E, right panel) detected a band of the same size in CBL as well as in *Hh*L. Interestingly, we detected GroEL in *Hh*L with serum IgG from DC-LMP1/CD40 animals with every age tested and this reactivity was increasing with the age of mice (Fig. 2F). In contrast, GroEL detection in *Hh*L with serum IgA from DC-LMP1/CD40 mice was observed only with sera obtained from mice at the age of 14 weeks and older (Fig. 2F). However, there was no GroEL-specific signal detected neither with serum IgG nor IgA from control mice (Fig. 2F), although their CBL did contain *Hh* (Fig. 2E and see below). Taken together, we identified the 60 kDa chaperonin GroEL from *Hh* as potential antigen recognized by the immune system during early colitis onset, indicating that *Hh* could be a disease driver in the DC-LMP1/CD40 colitis model.

### *Helicobacter hepaticus*-free DC-LMP1/CD40 mice are protected from early disease onset

The intestinal bacterium *Hh* is associated with IBD and induces spontaneous colitis in immunodeficient mice with severe combined immunodeficiency or IL10-deficient mice ^19, 21^. The fact, that we found *Hh*-specific Ig in sera of DC-LMP1/CD40 mice suggested that this mouse colony was endemically infected by *Hh*. To test this, we screened the fecal content from mice for presence of *Helicobacter* by genus-as well as species-specific PCR (Fig. 3A). We found the genus *Helicobacter* (*Hspp)* throughout all DC-LMP1/CD40 and control littermates (Fig. 3A). Moreover, all control littermates were consistently colonized with *Hh* (Fig. 3A, B). Surprisingly, young DC-LMP1/CD40 mice showed reduced prevalence already in week 3 of age, when only 57.1 % *Hh*-positive transgenic animals could be detected, in contrast to 100 % control littermates (Fig. 3B). Furthermore, *Hh* was hardly detectable in older DC-LMP1/CD40 mice, as in 10-week-old animals only 8.3 % were *Hh*-positive as compared to 100% of control littermates (Fig. 3B). Notably, we obtained similar results for colonization with *H. typhlonius* (*Ht*) (Fig. S4). In contrast, all animals tested were also colonized by *H. rodentium* (*Hr*), explaining consistent *Hspp* positive results (Fig. S4). None of the animals was tested positive for *H. bilis* (*Hb*) (Fig. S4). Taken together, conventionally-housed mice were endemically colonized with *Hh*. The fact, that DC-LMP1/CD40 animals show loss of *Hh* colonization in particular upon colitis progression suggests that these bacteria are eliminated by either ongoing immune responses or displacement by other bacteria during dysbiosis.

**Figure 3:**
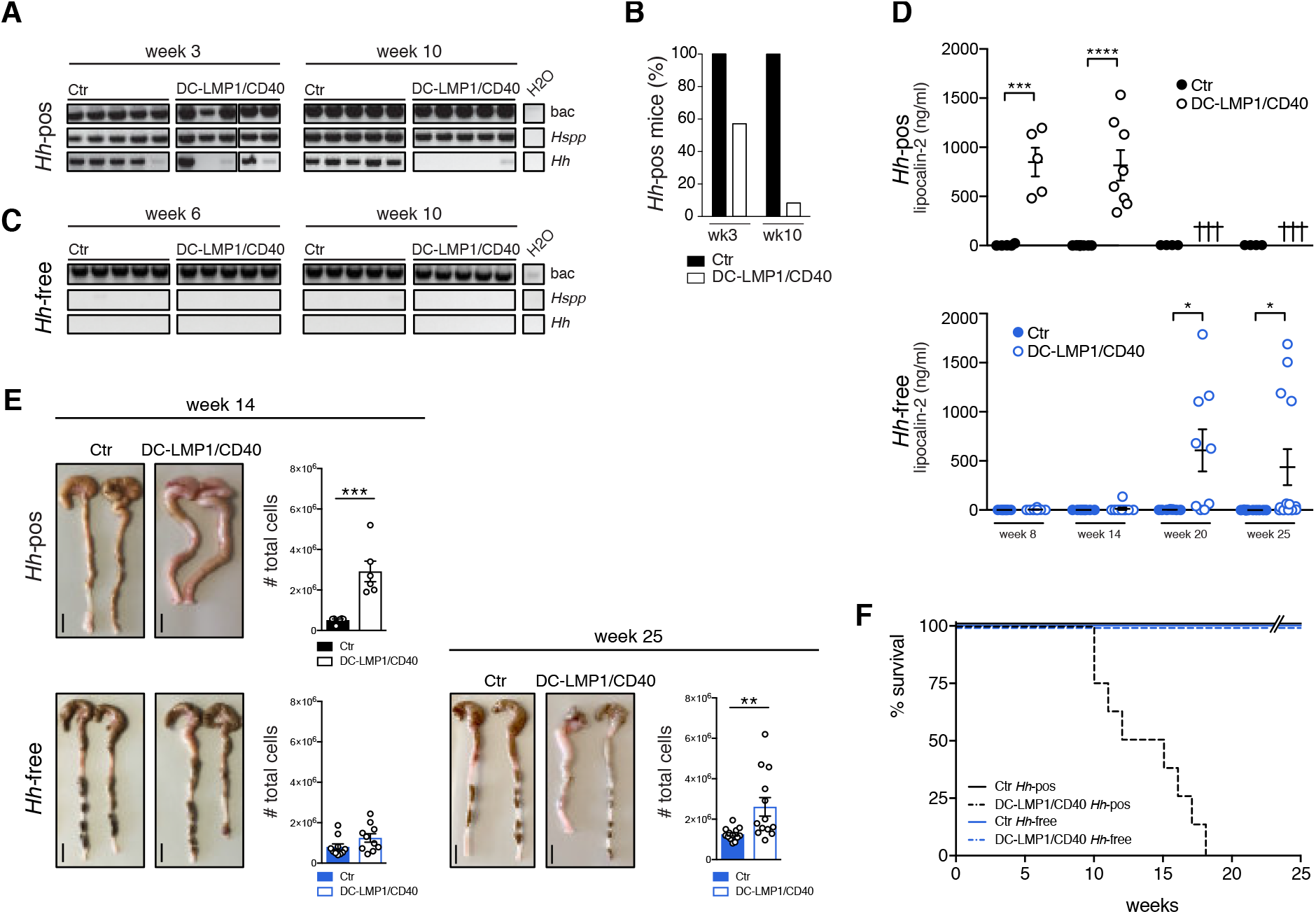
*Hh*-free DC-LMP1/CD40 are protected from early disease onset. (A) Bacterial DNA was extracted from Ctr or DC-LMP1/CD40 mice before rendering them *Hh*-free at the indicated time points. 16S rRNA gene primers were used to detect the species indicated and amplicons were analyzed by gel electrophoresis (n=7-14, shown are n=5 per group). (B) Data for *Hh*-pos animals from (A) is represented as bar graphs, illustrating the percentage of 3-or 10-week-old Ctr and DC-LMP1/CD40 animals tested positive for *Hh* before rendering them *Hh*-free (n=7-14 per group). (C) Bacterial DNA was extracted from Ctr or DC-LMP1/CD40 mice after rendering them *Hh*-free at the indicated time points. 16S rRNA gene primers were used to detect the species indicated and amplicons were analyzed by gel electrophoresis (n=5-9, shown are n=5 per group).(D) Levels of fecal lipocalin-2 were measured by ELISA in *Hh*-pos (upper panel) or *Hh*-free (lower panel) Ctr and DC-LMP1/CD40 mice at the indicated time points. Shown are data from two pooled experiments for *Hh*-pos animals (n=5-9) and for *Hh*-free animals (n=9-13) as mean ± SEM. Crosses represent already dead animals at the indicated time points. (E) Macroscopic pictures as well as total cell number of colons from *Hh*-pos (upper panel) or *Hh*-free (lower panel) Ctr and DC-LMP1/CD40 animals at the indicated time points. Shown are two representative colon pictures per group with scale bars = 1 cm. Bar graphs show total colon cell numbers in Ctr and DC-LMP1/CD40 mice from three pooled experiments (mean ± SEM, n=6-13). (F) Kaplan-Meier plot showing survival of *Hh*-free and *Hh*-pos Ctr and DC-LMP1/CD40 animals (n=10). Data for *Hh*-pos animals were taken from figure 2 in our previous publication ^38^ .bac: bacteria; *Hspp*: *Helicobacter species*; *Hh*: *H. hepaticus*

By embryo transfer we rederived mice to an *Hh*-free specific-pathogen-free (SPF) colony. *Hh-*colonization status was confirmed by genus- and species-specific PCR with fecal content from 6- and 10-week-old mice (Fig. 3C). Notably, all animals were tested negative for *Hh* (Fig. 3C) as well as *Ht*, *Hr* and *Hb* (Fig. S4). None of the *Hh*-free DC-LMP1/CD40 animals showed elevated fecal lipocalin-2 levels at the age of 8 or 14 weeks, when *Hh*-positive (*Hh*-pos) DC-LMP1/CD40 mice had already significantly elevated fecal lipocalin-2 levels (Fig. 3D). Interestingly, we did detect significantly increased fecal lipocalin levels only at much later time points in some but not all *Hh*-free DC-LMP1/CD40 mice (Fig. 3D). At week 20, 55.5 % and at week 25 30.8 % of *Hh*-free transgenic mice showed elevated lipocalin-2 levels (Fig. 3D). When we compared the phenotype of *Hh*-pos and *Hh*-free animals at the age of 14 weeks, we observed neither macroscopic signs of colitis, nor elevated total cell numbers in the colonic LP of *Hh*-free DC-LMP1/CD40 animals (Fig. 3E). In contrast, 14 weeks old *Hh*-pos DC-LMP1/CD40 mice already showed signs of colitis such as shortened and thickened colon as well as strong increase in total colonic cell numbers (Fig. 3E and ^24^). After 25 weeks, some *Hh*-free DC-LMP1/CD40 mice also showed an inflamed phenotype with a shortened and thickened colon as well as increased cell numbers infiltrating the colon LP (Fig. 3E). Of note, *Hh*-free DC-LMP1/CD40 mice not only showed no morbidity but also normal survival rates. Compared to *Hh*-positive DC-LMP1/CD40 animals, which usually die between 10 to 18 weeks of age (Fig. 3F, ^38^), none of *Hh*-free transgenic animals died before week 25, when they were finally analyzed (Fig. 3F). Our data shows a substantial delay in disease onset as well as less morbidity of *Hh*-free DC-LMP1/CD40 mice, indicating a crucial role for this microbe in disease initiation and outcome in CD40-mediated colitis.

### Conventionally-housed but not SPF-housed DC-LMP1/CD40 mice show changes in intestinal taxa composition upon colitis onset

To reveal the role of commensals in colitis initiation, we further analyzed intestinal taxa composition at family level in both conventionally-housed and SPF-housed mice by 16S rRNA gene sequencing of fecal samples (Fig. 4). In conventionally-housed mice, microbial changes in 8-week-old mice turned out to be genotype-dependent when compared to 3-week-old mice (Fig. 4A). We observed microbial changes in colitis-diseased 8-week-old DC-LMP1/CD40 animals when compared to control littermates, indicating dysbiosis upon colitis onset, confirming our previous finding of reduced microbial diversity in diseased transgenic mice ^24^ (Fig. 4A). As expected, when we rederived the mice to an *Hh*-free SPF microbiota, the overall complexity of taxa composition at family level was strongly reduced, independent of their genotype or age (Fig. 4A). Interestingly, we could also not observe substantial differences in the taxonomic profile within 20-week-old SPF-housed DC-LMP1/CD40 mice (Fig. 4A). The differential taxa abundance in 8-week-old conventionally-housed and 20-week-old SPF-housed mice was further determined using the analysis composition of microbiomes (ANCOM) function (Fig. 4B). Here, conventionally-housed DC-LMP1/CD40 mice showed *Enterobacteriaceae* blooming, characteristic for dysbiosis during colitis ^39, 40^, but also increased abundance of *Peptostreptococcacea*e, *Turicibacteraceae*, and *Enterococcaceae*, while we observed only increased abundance of *F16* in control littermates (Fig. 4B, upper panel). In contrast, both SPF-housed transgenic and control littermates showed a very homogeneous and strongly limited microbial complexity (Fig. 4B, lower panel). Taken together, *Hh*-free SPF transgenic and control mice show similar microbial composition and no dysbiosis is evident in aged *Hh*-free DC-LMP1/CD40 mice.

**Fig. 4:**
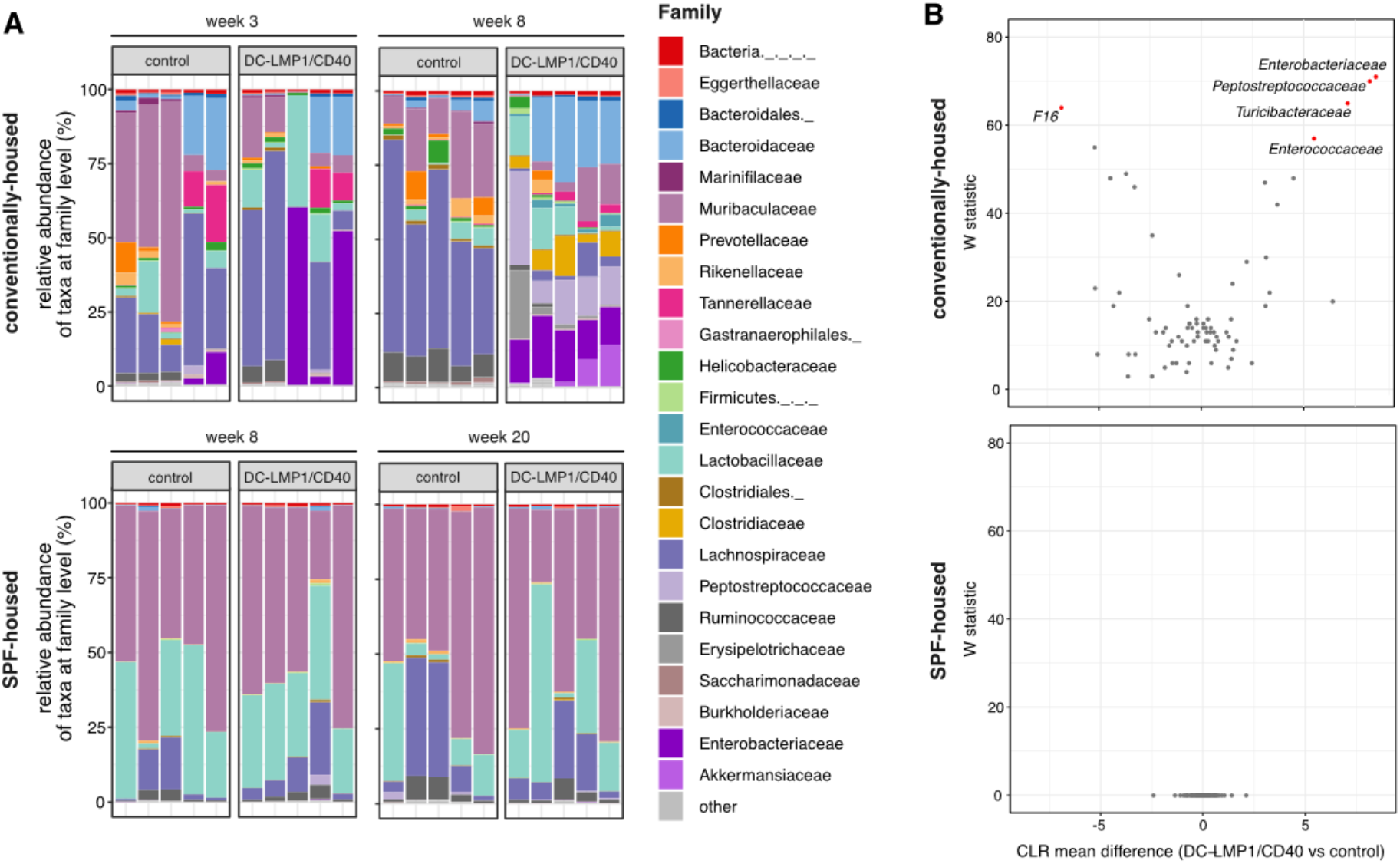
Conventionally-housed but not SPF-housed DC-LMP1/CD40 animals show changes in taxa composition. Analysis of the intestinal microbiota in fecal samples from conventionally-housed or SPF-housed control and DC-LMP1/CD40 mice at the indicated time points was based on sequencing the V3-V4 variable regions of the 16S rRNA gene (Illumina MiSeq). Filtered sequences were further processed using Qiime2 version 2020.2. A) Shown is the relative abundance of taxa at family level with each bar representing one animal (n=5 per group). Taxonomic assignment was performed with *classify-sklearn* using a classifier trained on SILVA database (Qiime version 132 99% 16S). B) Differential abundance in 8-week-old conventionally-housed or 20-week-old SPF-housed DC-LMP1/CD40 mice vs control littermates was estimated using the ANCOM function after collapsing to taxonomic level 5 and adding pseudo counts. CLR: Centered Log Ratio.

### DC-LMP1/CD40 mice rapidly develop strong intestinal inflammation upon colonization with *Hh*

To investigate if *Hh* is causal for disease initiation, we inoculated 8-week-old *Hh*-free animals with *Hh* (strain ATCC 51448) by oral gavage (Fig. 5A). Already at day 21 post inoculation (p.i.), all DC-LMP1/CD40 mice and control littermates, but not PBS-treated mice were *Hh*-positive as shown by species-specific PCR from feces (Fig. 5B). At day 40 p.i., when mice were finally sacrificed for analysis, all *Hh*-colonized mice were still *Hh* positive (Fig. 5B). Of note, all mice were negative for the other most relevant *Hspp* which are also routinely tested according to FELASA recommendations, confirming mono-colonization with *Hh* by oral gavage (Fig. S5). Furthermore, *Hh*-infected DC-LMP1/CD40 mice did show significantly elevated fecal lipocalin-2 levels compared to control littermates already on day 21 p.i., indicating a rapid disease onset upon colonization with *Hh* (Fig. 5C). By d40 p.i., *Hh*-infected DC-LMP1/CD40 mice, but not control littermates showed a strong increase in cells infiltrating the colonic LP as well as a shortened and thickened colon, indicating ongoing inflammation and colitis (Fig. 5D). In contrast, PBS-treated DC-LMP1/CD40 and control mice did not have elevated lipocalin-2 levels in their feces (Fig. 5C), neither did they show elevated cell numbers nor macroscopic changes of the large intestine (Fig. 5D) as observed previously in 14-week-old mice (Fig. 3D, E). Thus, our data reveal that *Hh* is rapidly provoking strong intestinal inflammation in DC-LMP1/CD40 mice, indicating that this bacterial stimulus combined with CD40-signaling on DCs and absent iTregs is causing the development of early onset colitis.

**Figure 5:**
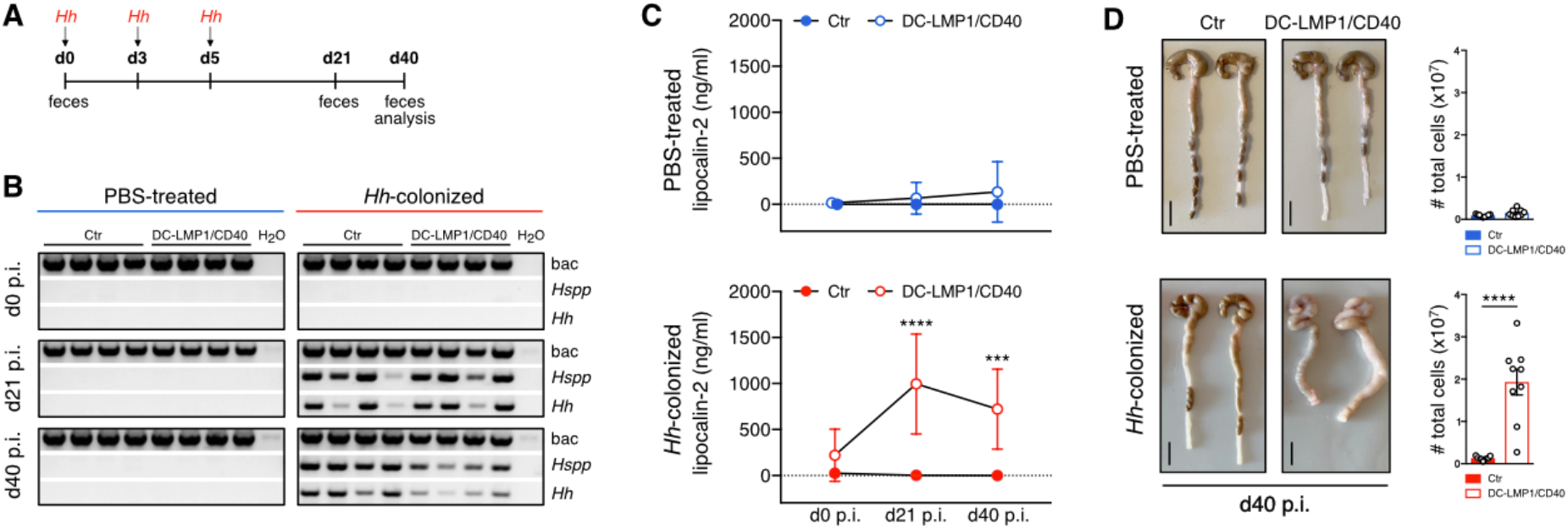
Strong intestinal inflammation upon *Hh*-recolonization. Schematic illustration of colonization of *Hh*-free Ctr and DC-LMP1/CD40 mice with a pure culture of *Hh* by oral gavage at the indicated time points. Feces were collected at the indicated time points and animals were sacrificed 40 days p.i. (B) Bacterial DNA was extracted from fecal samples at the indicated time points from PBS-treated or *Hh*- colonized Ctr and DC-LMP1/CD40 mice at the indicated time points. *Hh*- colonization was confirmed by PCR. Shown is one representative experiment out of two (n=4). (C) Fecal lipocalin-2 levels in PBS-treated or *Hh*-colonized Ctr and DC-LMP1/CD40 mice were determined by ELISA at the indicated time points. Data is shown as scatter plot for two pooled experiments with mean ± SEM (n=9). (D) PBS-treated or *Hh*-colonized Ctr and DC-LMP1/CD40 mice were sacrificed at day 40 p.i‥ Shown are macroscopic pictures of two representative colons per group (scale bars = 1 cm) as well as bar graphs, representing total colon LP cell numbers from two pooled experiments with mean ± SEM (n=9). Bac: bacteria; *Hspp*: *Helicobacter* species; *Hh*: *H. hepaticus*

### *Helicobacter hepaticus* affects colonic CD4^+^ T cell differentiation

We previously reported the effect of constitutive CD40-signaling on intestinal DCs ^24^. Transgenic animals did show a strong reduction of tolerogenic CD103^+^ DC subsets in the colonic LP and mLNs. As a consequence, RORγt^+^Helios^−^ iTreg generation was drastically impaired in the large intestine of DC-LMP1/CD40 animals. As the mouse colony used for our previous study was endemically infected by *Hh* (Fig. 3A, B), we wondered whether *Hh* is responsible for the phenotypical changes we observed in this colitis model. Therefore, we next analyzed cell subsets in the colonic LP of *Hh*-free DC-LMP1/CD40 and control littermates. We did find a strong reduction in CD103^+^CD11b^−^ as well as CD103^+^CD11b^+^ intestinal DCs in 14-week-old *Hh*-free DC-LMP1/CD40 animals but not control littermates (Fig. 6A), similar to what we previously described for *Hh*-pos animals ^24^. Furthermore, RORγt^+^Helios^−^ iTregs were also significantly reduced in the colonic LP of 14-week-old *Hh*-free DC-LMP1/CD40 mice, but not control animals (Fig. 6B) comparable with our previous findings in *Hh*-pos animals ^24^. Of note, the reduction of intestinal CD103^+^ DCs as well as RORγt^+^Helios^−^ iTregs was also found in 25-week-old *Hh*-free DC-LMP1/CD40 mice, but not control animals (Fig. S6). Therefore, we conclude that the phenotypical changes in DC-LMP1/CD40 mice are rather a consequence of the transgene expression in DCs, suggesting that *Hh* has no direct impact on DC or Treg differentiation in this model.

**Figure 6:**
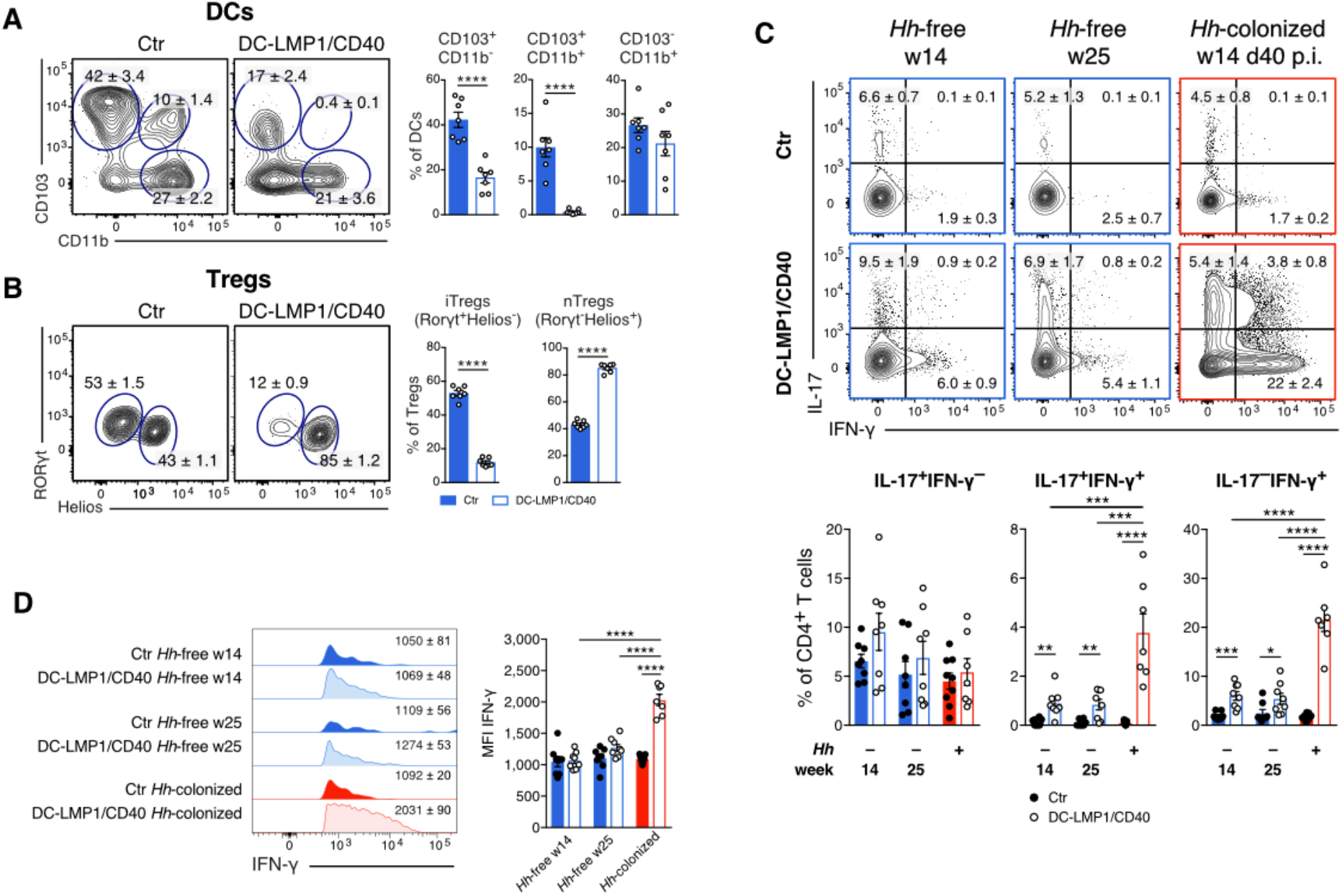
*Hh* affects colonic T cell-fate decisions. Different cell subsets in the colonic LP were analyzed in 14-week-old *Hh*-free Ctr and DC-LMP1/CD40 animals. Shown are representative FACS-plots as well as pooled statistics from two experiments (mean ± SEM, n=7), illustrating frequencies of the indicated cell subsets. (A) DCs were gated on single, live, CD45^+^, MHCII^+^CD11c^+^, CD64^−^ cells. (B) Tregs were gated on single, live, CD45^+^, CD3^+^CD4^+^, FoxP3^+^CD25^+^, RORγt^+^Helios^−^ (iTregs) or RORγt-Helios^+^ (nTregs). Single-cell suspensions of the colonic LP from 14- and 25-week-old *Hh*-free or *Hh*-infected Ctr or DC-LMP1/CD40 (14-weeks-old, 40 days post *Hh-*infection) mice were stimulated with PMA/Ionomycin and subsequently stained intracellularly for IL-17 and IFN-γ production at the indicated time points. Bar graphs represent pooled statistics from two experiments (mean ± SEM, n=7-9) animals. (C) T cells were pre-gated on single, live, CD45^+^, CD3^+^CD4^+^ cells. Shown are representative FACS-plots as well as bar graphs, illustrating the frequencies of indicated cell subsets within the CD4 T cell population. (D) Shown are representative histograms as well as bar graphs, illustrating the MFI of IFN-γ expression within cells within IFN-γ^+^ CD4^+^ T cells from (C) as median ± SEM.

We also know from our previous study that DC-LMP1/CD40 mice show a strong increase in IL-17^+^IFN-γ^+^ Th17/Th1 and IFN-γ^+^ Th1 cells in the colonic LP, indicating that non-pathogenic Th17 cells in DC-LMP1/CD40 mice are differentiating into pathogenic Th1 cells ^24^. To evaluate the role of *Hh* in this CD4^+^ T cell differentiation process, we compared IL-17- and IFN-γ-producing CD4^+^ T cells in different mice (Fig. 6C). We did not detect significantly different frequencies of IL-17^+^CD4^+^ T cells in DC-LMP1/CD40 or control animals, neither in *Hh*-free nor in *Hh*-infected animals (Fig. 6C). However, at day 40 p.i., *Hh*-infected DC-LMP1/CD40 mice had significantly increased frequencies of both, IL-17^+^IFN-γ^+^ Th17/Th1 and IFN-γ^+^ Th1 cells when compared to 14- or 25-week-old *Hh-*free DC-LMP1/CD40 mice (Fig. 6C). Of note, also *Hh*-free transgenic animals showed some induction of IL-17^+^IFN-γ^+^ Th17/Th1 and IFN-γ^+^ Th1 cells in the colonic LP when compared to appropriate control littermates (Fig. 6C). However, when we analyzed the mean fluorescence intensity (MFI) of IFN-γ expression in IFN-γ^+^ CD4^+^ T cells, only Th1 cells from *Hh*-infected DC-LMP1/CD40 mice produced significantly higher amounts of IFN-γ when compared to *Hh*-free transgenic animals (Fig. 6D). Taken together, we could show that *Hh* has the potential to rapidly initiate the transdifferentiation of non-pathogenic Th17 into pathogenic Th1 cells in the colonic LP devoid of tolerogenic CD103^+^ DCs and iTregs. However, this is not an exclusive property of *Hh* but can also be accomplished in *Hh*-free mice by other commensals or mechanisms, yet with much less efficacy.

## Discussion

In this study we identified the murine commensal *Helicobacter hepaticus* as driver of the pathogenesis in a CD40-mediated model of colitis. Upon very early colitis onset, DC-LMP1/CD40 animals showed elevated serum IgG- as well as IgA-levels. Although IgA is mainly produced locally in the gut, we observed elevated IgA levels and increased anti-commensal IgA also in sera of mice. To ensure mucosal homeostasis, the gut sustains tolerance towards commensal bacteria by restraining them by various mechanisms, including the secretion of protective anti-microbial peptides and bacteria-specific IgA. Thus, bacteria-specific antibodies are not detectable in sera from healthy SPF-housed mice ^41^. However, during inflammatory conditions systemic antibodies can be produced as a consequence of mucosal barrier dysfunction and thus increased exposure of commensals to systemic sites ^42^. The presence of commensal-specific serum antibodies in DC-LMP1/CD40 mice therefore suggests that compartmentalization might be broken in mice with colitis, leading to systemic antibody responses.

Only recently, it was reported that especially members of Proteobacteria are able to induce T cell-dependent serum IgA responses in conventionally-housed mice to protect them from lethal sepsis ^43^. In this study commensal *Helicobacter muridarum* was identified as driving species, which would induce mucosal IgA-secreting plasma cells as well as IgA^+^ bone marrow plasma cells ^43^. Our data suggests that dysbiosis in DC-LMP1/CD40 mice affects dissemination of bacteria, inducing systemic IgG as well as IgA production. This hypothesis is also supported by human studies, reporting elevated serum antibody levels in IBD patients ^44, 45^. However, we also found certain levels of bacteria-specific serum IgG in control littermates. One explanation for this observation might be that we used conventionally- but not SPF-housed mice for parts of our study. This seems more analogous to healthy humans, where also some level of systemic bacteria-specific IgG has been reported, which do increase further during IBD ^42^.

We identified serum antibodies from transgenic DC-LMP1/CD40 animals being bacteria-specific and recognizing a 60 kDa protein from *Hh*. The fact that only the 60 kDa chaperonin from *Hh* was identified with this method was surprising, but heat shock proteins have been reported as immunodominant antigens, inducing humoral and cellular immune responses to several diseases in humans and mice. For instance, αHsp60 antibodies are found in patients with tuberculosis and in mice infected with *Mycobacterium tuberculosis* ^46, 47^. Pathogen-derived 60 kDa chaperonin induces pro-inflammatory cytokines *in vitro* ^48^ and mice infected with *Yersinia enterocolitica* produce 60 kDa chaperonin-specific T cells involved in anti-pathogenic immune response ^46^. Serum antibodies specific for *H. pylori* Hsp60 were also reported in patients with gastric cancer ^49^. Our data suggest that *Hh* is involved in disease development and its 60 kDa chaperonin might be an immunodominant antigen in the CD40-mediated colitis model. *Hh* is known as pathobiont, endemic in many mouse colonies ^15, 16^ where it can elicit intestinal inflammation in immunodeficient or immunocompromized mice. This mimicks human IBD as demonstrated by several mouse models where it elicits spontaneous colitis ^19, 20, 21, 23, 50^. Although all of our conventionally-housed mice were tested positive for the *Helicobacter* genus, only control mice remained consistently positive for *Hh*. In contrast, young DC-LMP1/CD40 mice showed already reduced prevalence while *Hh* was hardly detectable in older DC-LMP1/CD40 mice. One reason for this phenomenon might be the clearance of *Hh*, eventually as a consequence of increased anti-*Hh* serum IgG and IgA levels. Alternatively, *Hh* may be simply displaced for example by *Enterobacteriaceae* which bloom ^39, 40^ during inflammation in DC-LMP1/CD40 animals. This may suggest that *Hh* is causing disease initiation but not its maintenance and progression.

In contrast, *Hh*-free DC-LMP1/CD40 mice only developed mild intestinal inflammation at the age of 5 to 6 months, when *Hh*-positive transgenic animals had already died from the disease. Interestingly, also IL-10-deficient mice developed intestinal inflammation with delayed onset and less severity in 5- to 6-month-old animals when maintained under SPF conditions ^21^. Also, *Hh*-free transgenic mice with Treg-specific c-Maf deficiency developed mild spontaneous colitis at a later age of 6 to 12 month ^50^. The protection from early disease onset in *Hh*-free DC-LMP1/CD40 mice suggests that *Hh* might be a very potent disease driver. Nevertheless, although we could not find any evidence for specific commensals involved in disease initiation in aged *Hh*-free SPF-housed transgenic mice, also other bacteria may cause disease, although much weaker, with later onset and in less mice.

While we determined *Hh* as disease driver in CD40-mediated colitis model, this microbe did not have a direct impact, neither on DC nor on Treg differentiation in the colon LP, as CD103^+^ DCs and RORγt^+^Helios^−^ iTregs were similarly reduced in both, *Hh*-free and *Hh*-infected DC-LMP1/CD40 animals ^24^. In contrast, CD4^+^ effector T cell differentiation in the colon LP was affected by *Hh*, which significantly increased IL-17^+^IFN-γ^+^ Th17/Th1 and IFN-γ^+^ Th1 cells. Also in IL-10^−/−^ mice *Hh*-infection induced pathogenic, *Hh*-specific IL-17^+^IFN-γ^+^ Th17/Th1 cells ^51^, probably due to the inability of Tregs to restrain colitogenic Th17 cells in *Hh*-positive IL-10^−/−^ mice ^50^.

Our data revealed that the intestinal microbiota is able to modulate the host immune response with impact on disease onset, progression and severity. Here, we identified *Hh* as disease driver in the DC-LMP1/CD40 colitis model. In the context of constitutive CD40-signaling in DCs, we could show that *Hh* induces early onset of fatal colitis, by causing the transdifferentiation of non-pathogenic Th17 cells into pathogenic Th1 cells in the colonic LP. Our results are also of relevance for other studies using conventionally-housed mice as *Hh* is endemic in many mouse colonies. Our data further confirm the important role of the gut microbial composition during health and disease and reveal that single bacterial species can dramatically affect host immunity. The identification of other potential disease driving bacteria as well as specific bacterial antigens and underlaying mechanisms in IBD is central. This further contributes to understanding the complex interaction of microbiota and host immune cells to develop and improve in particular personalized therapeutic strategies in IBD.

## Material and Methods

### Mice

DC-LMP1/CD40 mice were generated as previously described ^24^. Briefly, CD11cCre mice ^52^ were crossed with LMP1/CD40^fl/flSTOP 53^ animals to obtain DC-LMP1/CD40 mice with constitutive CD11c-specific CD40-signalling.

Mice were analyzed in sex- and age-matched groups of 8 - 25 weeks of age, unless otherwise stated. Littermate animals were used as controls in a non-randomized, non-blinded fashion. Animal experiment permissions were granted by the animal ethics committee Regierung von Oberbayern, Munich, Germany (55.2.1.54-2532-22-2017). Mice were bred and maintained under conventional conditions at the animal facility of the Institute for Immunology, Ludwig-Maximilians-Universität München. After embryo transfer rederivation performed by ENVIGO (Huntingdon, United Kingdom), all mice were kept under specified pathogen-free conditions (tested quarterly according to FELASA-14 recommendations) and housed in groups of 2-3 animals in IVCs (Tecniplast, Germany) at a 12h/12h light/dark cycle. Mice had free access to water (acidified and desalinated) and standard rodent chow (Altromin, 1310M).

### Single-cell preparation

Single-cell suspensions of lymph nodes were prepared by mashing organs through a 100 μM cell strainer. Samples were washed with PBS and stored on ice for further analysis. Number of living cells was determined using the CASY Counter (OMNI Life Science). Cells from the colonic LP were isolated as previously described ^24^. Briefly, the colon was removed, cleaned from fecal content, opened longitudinally, cut into pieces and predigested in Hank’s balanced salt solution (HBSS) supplemented with 10 mM HEPES and 10 mM EDTA for 10 min on a shaker at 37 °C. Pieces were further digested for 30 min and then twice for 20 min with a mixture of Collagenase IV (157 Wuensch units ml^−1^, Worthington), DNAse I (0.2 mg ml^−1^ dissolved in PBS) and Liberase (0.65 Wuensch units ml^−1^, both Roche, dissolved in HBSS supplemented with 8 % FCS). Lymphocytes were purified with a 40/80 Percoll gradient and the number of living cells was determined using the CASY Counter.

### Flow cytometry analysis

Where possible, 2 × 10^6^ cells were stained with titred antibodies in PBS containing 2 % FCS and 0.01 % NaN_3_ (FACS buffer) for 20 min at 4 °C in the dark. Cells were washed once and used for direct acquisition on BD FACSCanto or fixed using 2 % paraformaldehyde in FACS buffer and measured the next day. Dead cells were excluded using Zombie Aqua Fixable Viability Kit (BioLegend, Cat: 423102). For intracellular cytokine stainings, cells were fixed and permeabilized for 30 min at 4 °C in the dark after extracelluar stainings using BD Cytofix/Cytoperm (Fixation and Permeabilization Solution, BD Biosciences, Cat: 51-2090KZ) according to manufacturer’s instructions. Cells were washed and stained with indicated antibodies in 50 μl BD Perm/Wash (Buffer, BD Biosciences, Cat: 51-2091KZ) for 30 min at 4 °C in the dark. For transcription factor staining, cells were fixed and permeabilized after extracellular stainings in 1x Fixation/Permeabilization solution (eBioscience, Cat: 00-5523-00) for 30 min at 4 °C in the dark according to manufacturer’s instructions. Cells were washed twice with 1x Permeabilization Buffer (eBioscience, Cat: 00-5523-00) and stained with the indicated antibodies in 50 μl 1x Permeabilization Buffer for 30 min at 4 °C in the dark. Afterwards, cells were washed once and acquired on BD FACSCanto.

The following antibodies were used: FoxP3 (FJK-16s; eFlour660, dil. 1:50), Helios (22F6; FITC, dil. 1:400), RORγt (AFKJS-9; PE, dil. 1:100) (eBioscience); CD25 (PC61; PerCP, dil. 1:400), CD103 (M290; PE, dil. 1:150) (BD Pharmingen); CD11b (M1/70; APC-eFluor780, dil. 1:400) (Invitrogen); CD3 (17A2; AlexaFluor488, dil. 1:400; Pe-Cy7, dil. 1:400), CD4 (RM4-5; PerCP, 1:800; GK1.5; APC-Cy7, dil. 1:400), CD11c (N418; Pe-Cy7, dil. 1:400), CD45 (30-F11; BV421, dil. 1:400), CD64 (X54-5/7.1; APC, dil. 1:200), IL-17A (TC11-18H10.1; PE, dil. 1:200), IFN-γ (XMG1.2; APC, dil. 1:400), MHC class II (I-A/I-E) (M5/114.15.2; FITC, PerCP, dil. 1:800) (BioLegend). Data analysis was performed using FlowJo version 10 (TreeStar, Ashland, OR, USA).

### Ex vivo T cell restimulation

2 × 10^6^ cells were stimulated for 4 h at 23°C with 40 ng ml^−1^ PMA and 1 μg ml^−1^ ionomycin in the presence of 2 μM Monensin (Golgi-Stop, BD Biosciences, Cat: 51-2092KZ). Cells were washed twice with FACS buffer and stained for extracellular markers, fixed/permeabilized and stained for intracellular markers as described above.

### ELISA for fecal lipocalin-2

Fecal samples were reconstituted in PBS containing 0.1% Tween 20 (100 mg ml^−1^) and vortexed for 20 min for homogenisation. Upon centrifugation for 15 min at 100 x g at 4 °C, supernatants were centrifuged again for 10 min at 10,000 x g at 4 °C. The supernatants were analyzed for lipocalin-2 content using Quantikine ELISA kit for mouse Lipocalin-2/NGAL (R&D Systems, Cat: MLCN20).

### Determination of serum antibody concentrations

Blood from mice was collected by terminal cardiac puncture and transferred into a Microtainer tube (BD Biosciences, Cat: 365963). After incubation at room temperature for at least 3 h, the coagulated blood was centrifuged at 8000 rpm for 5 min at 21 °C and serum was frozen at −20°C until use. Serum antibody concentrations were determined using Mouse IgG total Ready-SET-Go! or Mouse IgA Ready-SET-Go! ELISA (eBioscience, Cat: 88-50400 and 88-50450), according to manufacturer’s instructions.

### Immunoprecipitation of bacterial antigens

Identification of bacterial antigens within the cecal bacterial lysate (CBL) was performed by using serum antibodies from control and DC-LMP1/CD40 mice for immunoprecipitation followed by Mass Spectrometry. Therefore, 50 μl protein G beads (Dynabeads Protein G, Invitrogen, Cat: 10004D) were coupled with 2.5 μg serum IgG from Ctr or DC-LMP1/CD40 mice for 10 min at room temperature. 1600 μg CBL was added to the coated beads for 30 min at room temperature and the complex was washed three times with PBS/Tween 0.02 % followed by additional 3 rounds of washing with 50 mM NH_4_HCO_3_. Samples were stored at −20°C until LC-MS/MS was performed by the Protein Analysis Unit (Biomedical Center, LMU Munich).

### On-beads trypsin digest and Mass Spectrometry

Following the immunoprecipitation procedure described above, beads were incubated with 100 μl of 10 ng μl^−1^ trypsin solution in 1 M Urea and 50 mM NH_4_HCO_3_ for 30 min at 25°C for trypsin digestion. The supernatant was collected, beads washed twice with 50 mM NH_4_HCO_3_ and all three supernatants collected together and incubated overnight at 25°C at 800 rpm after addition of dithiothreitol to 1 mM. Iodoacetamide was added to a final concentration of 27 mM and samples were incubated at 25°C for 30 min in the dark. 1 μl of 1 M dithiothreitol was added to the samples and incubated for 10 min to quench the iodoacetamide. Finally, 2.5 μl of trifluoroacetic acid was added and the samples were subsequently desalted using C18 Stage tips. Samples were evaporated to dryness, resuspended in 15 μl of 0.1 % formic acid solution and injected in an Ultimate 3000 RSLCnano system (Thermo), separated in a 15-cm analytical column (75 μm ID home-packed with ReproSil-Pur C18-AQ 2.4 μm from Dr. Maisch) with a 50 min gradient from 5 to 60 % acetonitrile in 0.1 % formic acid. The effluent from the HPLC was directly electrosprayed into a Qexactive HF (Thermo) operated in data dependent mode to automatically switch between full scan MS and MS/MS acquisition. Survey full scan MS spectra (from m/z 375 - 1600) were acquired with resolution R = 60,000 at m/z 400 (AGC target of 3 × 10^6^). The 10 most intense peptide ions with charge states between 2 and 5 were sequentially isolated to a target value of 1 × 10^5^, and fragmented at 27 % normalized collision energy. Typical mass spectrometric conditions were: spray voltage, 1.5 kV; no sheath and auxiliary gas flow; heated capillary temperature, 250°C; ion selection threshold, 33,000 counts. MaxQuant 1.5.2.8 was used to identify proteins and quantify by intensity-based absolute quantification (iBAQ) with the following parameters: Database, uniprot_proteomes_Bacteria_151113.fasta; MS tol, 10 ppm; MS/MS tol, 10 ppm; Peptide FDR, 0.1; Protein FDR, 0.01 Min. peptide Length, 5; Variable modifications, Oxidation (M); Fixed modifications, Carbamidomethyl (C); Peptides for protein quantitation, razor and unique; Min. peptides, 1; Min. ratio count, 2. Identified proteins were considered as interaction partners if their MaxQuant iBAQ values were greater than log2 2-fold enrichment and p-value 0.05 (ANOVA) when compared to the control. The mass spectrometry proteomics data have been deposited to the ProteomeXchange Consortium via the PRIDE (http://www.proteomexchange.org) partner repository with the dataset identifier PXD018025.

### Culture and lysate preparation of *Hh*

The *Helicobacter hepaticus* strain *Hh*-2 (ATCC 51448) ^54^ was purchased from the Leibniz Institute DSMZ - German Collection of Microorganisms and Cell Cultures (DSM No.22909) and cultivated at the Max von Pettenkofer-Institute, LMU Munich. Bacteria from cryo stock were resuspended in Brain Heart Infusion (BHI) medium and put onto blood agar plates (Columbia agar with 5 % sheep blood, BD, Cat: 4354005). Plates were incubated in a chamber with anaerobic conditions (83 % N_2_, 10 % CO_2_, 7 % H_2_) for 4 days at 37 °C. A subculture was cultivated further on in BHI medium with 3 % sheep serum in a culture flask in the chamber with anaerobic conditions for additional 4 days at 37 °C.

For *Hh* lysate (*Hh*L) preparation, bacterial cells were harvested and washed 2 - 3 times with PBS. Cell pellets were resuspended in PBS and lyzed by sonification with the Sonifier 150 Cell Disruptor (Branson) 6 times for 3 min at level 3 on ice. Lyzed cells were centrifuged at 20,000 x g for 30 min at 4°C and the supernatant was mixed with protease inhibitor (cOmplete ULTRA Tablets, Roche, Sigma-Aldrich, Cat: 05892953001). Protein concentration was determined using the Qubit Protein Assay Kit and Fluorometer (Invitrogen), according to the manufacturer’s instructions and the lysate was stored at −20 °C until use for immunoblot or ELISA.

### ELISA for commensal- or *Hh*-specific antibodies

This assay was performed as previously described ^24^ with the following modifications. The CBL was diluted in carbonate buffer to a final concentration of 1 μg ml^−1^. *Hh*L was prepared as described above and diluted in carbonate buffer to a final concentration of 0.1 μg ml^−1^. Differences in serum antibody concentrations between Ctr and DC-LMP1/CD40 mice were adjusted by using 2.5 μg ml^−1^ serum IgG or 6.5 μg ml^−1^ serum IgA for all samples.

### Immunoblotting for commensal- or *Hh*-specific antibodies

Serum IgG or IgA reactivity towards CBL or *Hh*L was analyzed by immunoblot analysis. 30 μg CBL or 20 μg *Hh*L were separated by SDS-PAGE and transferred to a nitrocellulose membrane. Sera of mice were used as primary antibodies. Differences in serum antibody concentrations between Ctr and DC-LMP1/CD40 mice were adjusted by using 2.5 μg ml^−1^ serum IgG or 1 μg ml^−1^ serum IgA for all samples. In some experiments, mouse IgG1 anti-human heat shock protein 60 (aHSP60) antibody (clone LK-2, Enzo) was additionally used as primary antibody (1:10,000 in PBS/1 % nonfat dried milk). HRP-conjugated secondary antibodies were used as follows: goat anti-mouse IgG-HRP (SouthernBiotech, Cat: 1030-05; 1:10,000) or goat anti-mouse IgA-HRP (SouthernBiotech, Cat: 1040-05; 1:10,000). Western Lightning Plus-ECL Detection Reagent (PerkinElmer) and X-ray films (Amersham) were used for protein detection.

### Bacteria screening PCR

Mice were screened for bacterial colonization by PCR using the 16S rRNA gene as target. Genomic DNA was isolated from fecal pellets with the QIAamp Fast DNA Stool Mini Kit (Qiagen), according to manufacturer’s instructions. 5 - 10 ng DNA was used for amplification with MyTaq Polymerase (Bioline). The PCR cycling conditions were as follows: denaturation at 94°C for 1 min, annealing for bacteria at 58°C, for *Hspp* and *Hh* at 61°C and for *Ht, Hr* and *Hb* at 55°C for 1 min, elongation at 72°C for 1 min (35 cycles) and final elongation at 72°C for 7 min.

The following primer sets were used: bacteria (forward primer: 5’-TCCTACGGGAGGCAGCAGT-3’, reverse primer: 5’-GGACTACCAGGGTATCTAATCCTGTT-3’, 467 bp) ^55^; *Hspp* (forward primer: 5’-TATGACGGGTATCCGGC-3’, reverse primer: 5’-ATTCCACCTACCTCTCCCA-3’, 375 bp) ^56^; *Hh* (forward primer: 5’-GCATTTGAAACTGTTACTCTG-3’, reverse primer: 5’-CTGTTTTCAAGCTCC- CC-3’, 417 bp) ^15^; *Ht* (forward primer: 5’-TTAAA-GATATTCTAGGGGTATAT-3’, reverse primer: 5’-TCTCCCATCTCTAGAGTGA-3’, 455 bp) ^57^; *Hr* (forward primer: 5’-GTCCTTAGTTGCTAACTATT-3’, reverse primer: 5’-AGATTTGCTCCATTTCACAA-3’, 166 bp) ^58^; *Hb* (forward primer: 5’-AGAACTGCATTTGAAACTACTTT-3’, reverse primer: 5’-GGTATTGCATCTCTTTGTATGT-3’, 638 bp) ^59^.

### 16S rRNA gene amplicon sequencing and taxonomic profiling

Microbiome analysis was done from whole DNA extracted from mouse fecal samples and is based on sequencing the V3-V4 variable regions of the 16S rRNA gene as previously described ^24^. Amplicons were analysed with mothur v. 1.43.0. (Schloss et al 75(23):7537-41) to remove chimeric sequences with the “chimera.vsearch”-command (default settings). Sequences were further processed using Qiime2 version 2020.2 Taxonomic assignment was performed with classify sklearn using a classifier trained on SILVA database (Qiime version 132 99 % 16S). Differential abundance was estimated using the ANCOM function ^60^ after collapsing to taxonomic level five and adding pseudo counts. 16S rRNA amplicon sequencing data have been deposited in the NCBI Sequence Read Archive under Accession Number SRX1799186.

### Colonization with *Hh* by oral gavage

Bacterial suspensions cultured as described above were used for oral inoculation to mice. *Hh* identity was confirmed by 16S RNA gene sequencing. Bacterial density was determined by OD measurements at 600 nm. Appropriate amount of suspension was washed with PBS and then adjusted to OD (600) = 3.0. 8-week-old mice were kept under specific and opportunistic pathogen free conditions for the time of the experiment and inoculated with 100 μl of the suspension by oral gavage at day 0, 3 and 5, for a total of 3 doses. Animals were analyzed 40 days post inoculation.

### Statistics

For absolute cell numbers, the percentage of living cells of a certain subset was multiplied by the number of living cells as determined by CASY Counter. Unless otherwise stated, significance was determined using unpaired Student’s *t*-test and defined as follows: *P<0.05, **P<0.01, and ***P<0.001 and ****P<0.0001. Error bars represent mean ± SEM.

## Acknowledgements

This work was supported by the Deutsche Forschungsgemeinschaft SFB1054 B03 to T.B. and A06 to A.K.; B.S., L.J. and D.R. were supported by the Center for Gastrointestinal Microbiome Research (CEGIMIR) of the German Center for Infection Research (DZIF). B.S. was supported by the DFG Priority Programme SPP1656 and the SFB1371. V.F. was supported by QBM, Munich.

## Author contributions

V.F. conducted the experiments. T.S. conducted bioinformatic analyses. S.S., L.J. and B.S. analysed sequences, D.R. cultivated Hh, I.F. and A.I. performed proteome analyses, A.K. B.P. and D.M. planned and performed colonization experiments with H.h., T.B. designed the experiments and V.F. and T.B. wrote the paper.

