## Supplemental data for "*Helicobacter hepaticus* as disease driver in a novel CD40-mediated model of colitis"

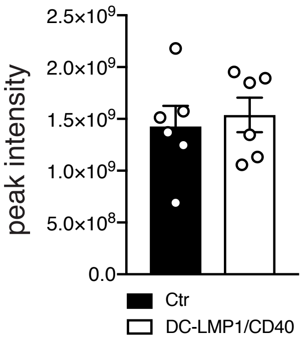


**Figure Suppl. 1: Intensities of Ig-related proteins (related to Fig. 2A).** Experiment was performed as in Fig. 2A, but data was filtered for Ig-related proteins only. Intensities of all Ig-related proteins detected by LC-MS/MS within every single control or DC-LMP1/CD40 sample are illustrated by bar graphs as mean ± SEM (two pooled experiments, n=6).

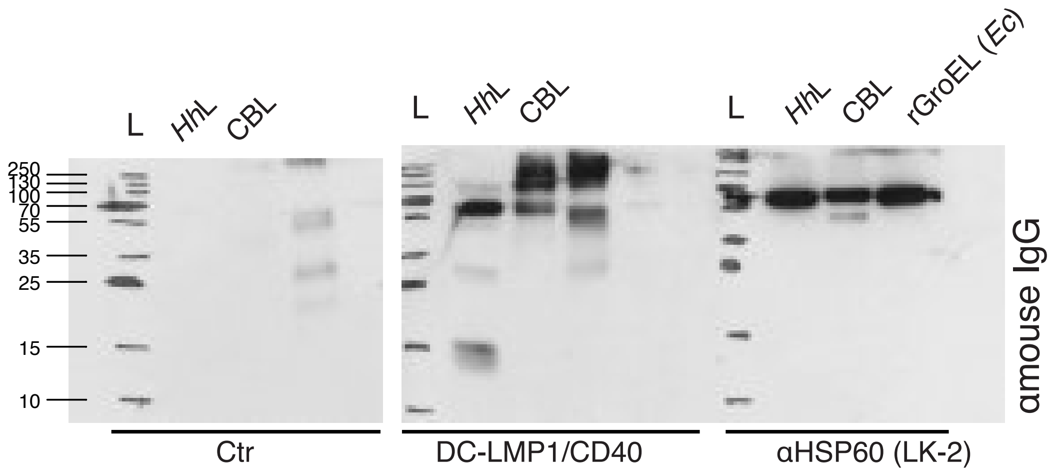


**Figure Suppl. 2: Full length Western Blot for the detection of a 60 kDa protein from *Hh*, related to Fig. 2E.** Full length blot for the detection of a 60 kDa protein from *Hh* displayed in Fig. 2E. 20 µg *Hh*L, 50 µg CBL or 0.5 µg recombinant GroEL from E. coli (rGroEL (Ec)) were separated by SDS-PAGE. Sera from one Ctr (left), one DC-LMP1/CD40 (middle) mouse and anti-HSP60 (clone LK-2, mouse IgG1 isotype) were used as primary antibodies. Anti-HSP60, immunogenic for recombinant human HSP60 was tested for cross-reactivity with the bacterial homolog GroEL, the rGroEL (Ec) and the 60 kDa chaperonin GroEL from *Hh*. Differences in serum antibody concentrations between Ctr and DC-LMP1/CD40 mice were adjusted by using same serum antibody concentrations (2.5 µg/ml IgG) as calculated from Fig. 1B. Anti-mouse IgG-HRP was used as secondary antibody. L: Ladder for 10-250 kDa.

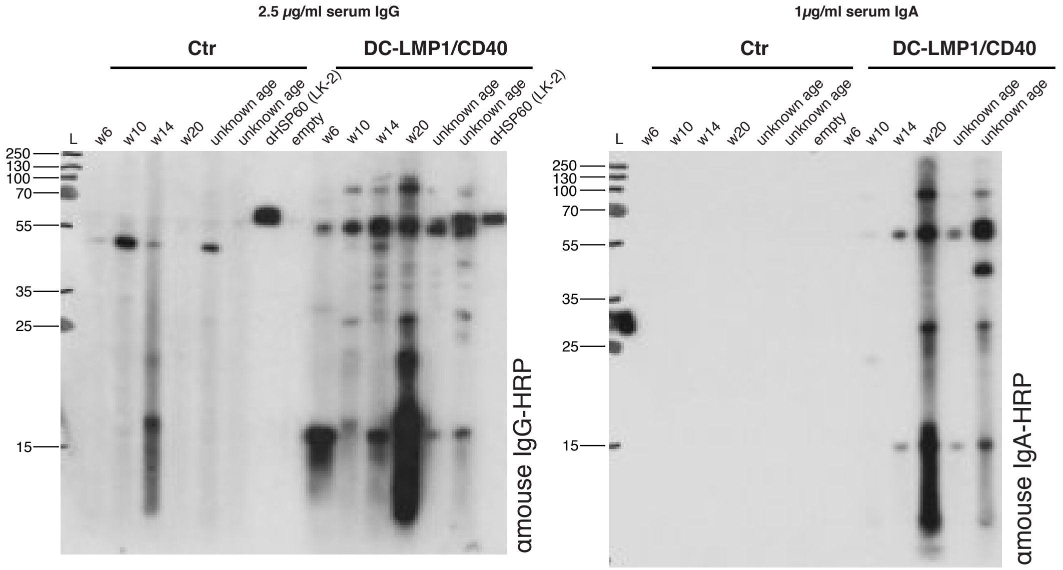


**Figure Suppl. 3: Full length Western Blot for serum screening, related to Fig. 2F.** Full length blot for sera-screening displayed in Fig. 2F. Detection of a 60 kDa protein from *Hh* by immunoblotting with serum from Ctr or DC-LMP1/CD40 mice. 200 µg *Hh*L was seperated by SDS-PAGE. Sera from Ctr or DC-LMP1/CD40 mice at the indicated age were screened with each lane representing one serum sample. Anti-HSP60 (clone LK-2, mouse IgG1 isotype) was used as primary antibody to detect the bacterial homolog of human HSP60, the 60 kDa chaperonin GroEL. Differences in serum antibody concentrations between Ctr and DC-LMP1/CD40 mice were adjusted by using same serum antibody concentrations (2.5 µg/ml IgG or 1 µg/ml IgA as calculated from Fig. 1B). Anti-mouse IgG-HRP (left) or anti-mouse IgA-HRP (right) were used as secondary antibodies. L: Ladder for 15-250 kDa.

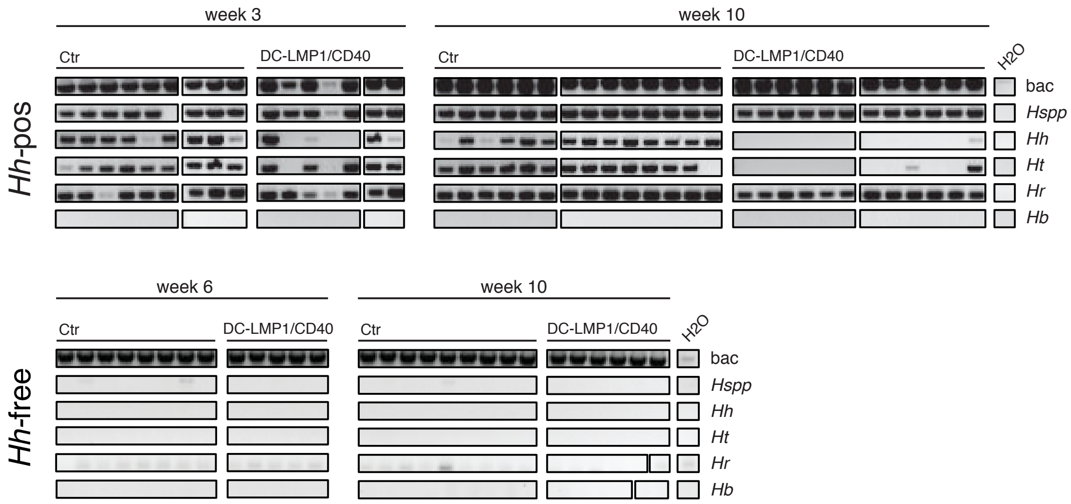


**Figure Suppl. 4: Screening of mice for colonization with *Helicobacter*, related to Fig. 3A, C.** Bacterial DNA was extracted from fecal samples from Ctr or DC-LMP1/CD40 mice before (*Hh*-pos, upper panel) and after rendering them *Hh*-free (lower panel) at the indicated time points. 16S rRNA gene primers were used to detect the species indicated and amplicons were analyzed by agarose gel electrophoresis (n=5-14). bac: universal bacteria; *Hspp: Helicobacter* species; *Hh*: *H. hepaticus*; *Ht*: *H. typhlonius*; *Hr*: *H. rodentium*; *Hb*: *H.bilis*

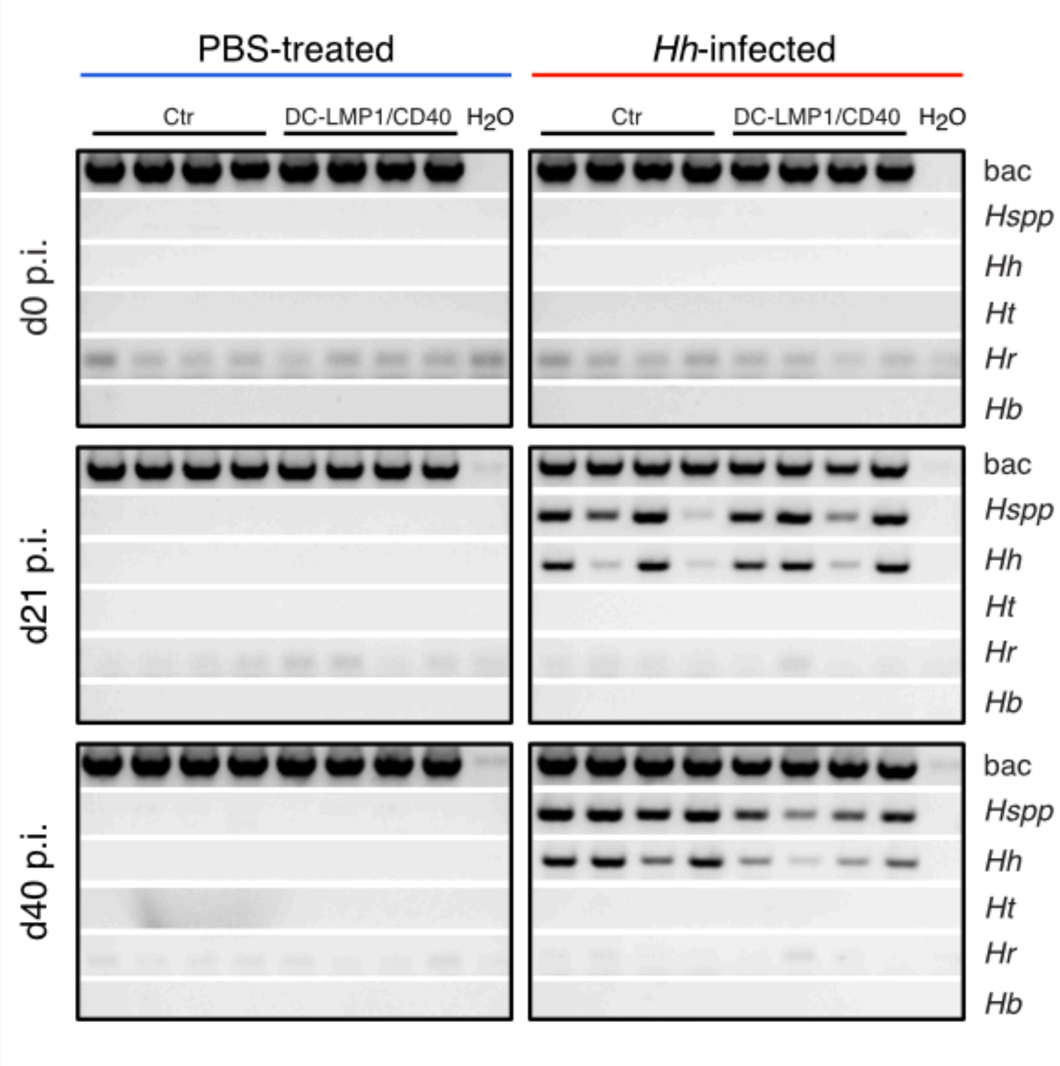


**Figure Suppl. 5: Screening of mice for colonization with *Helicobacter* upon infection, related to Fig. 5B.** Bacterial DNA was extracted from fecal samples from PBS-treated Ctr or DC-LMP1/CD40 mice and from animals inoculated with *Hh* at the indicated time points. 16S rRNA gene primers were used to detect the species indicated and amplicons were analyzed by agarose gel electrophoresis. Shown is one representative experiment out of two (n=4). bac: universal bacteria; *Hspp*: *Helicobacter* species; *Hh*: *H. hepaticus*; *Ht*: *H. typhlonius*; *Hr*: *H. rodentium*; *Hb*: *H. bilis*

**Commented [VF1]:** Hier auch Hh-colonized statt -infected?! Hab ich im affinity file abgeändert

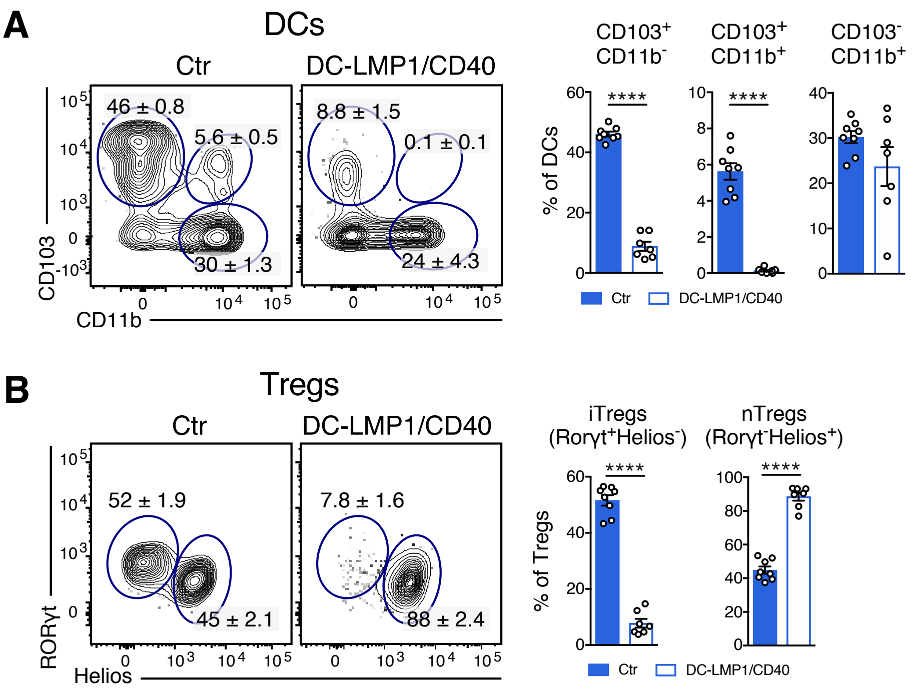


**Figure Suppl. 6: Loss of intestinal CD103$^+$ DCs and iTregs in 25-week-old *Hh*-free DC-LMP1/CD40 mice. Related to Fig. 6A, B.** (A-B) Different cell subsets in the colonic LP were analyzed in 25-week-old *Hh*-free Ctr and DC-LMP1/CD40 animals. Shown are representative FACS-plots as well as pooled statistics from two experiments (mean ± SEM, n=7-8), illustrating frequencies of the indicated cell subsets.

(A) DCs were gated on single, live, CD45$^+$, MHCII$^+$CD11c$^+$, CD64$^-$ cells.

(B) Tregs were gated on single, live, CD45$^+$, CD3$^+$CD4$^+$, FoxP3$^+$CD25$^+$, RORγt$^+$Helios (iTregs) or RORγtHelios$^+$ (nTregs).
